# Nicotine-driven hyperactivation of larval locomotion

**DOI:** 10.64898/2026.01.20.700145

**Authors:** Stephanie Dancausse, Jocelyn Robles, Jenna Fitzpatrick, Carolyn Garcia, Oshani Fernando, Anastasiia Evans, James D. Baker, Mason Klein

## Abstract

Balance between excitatory and inhibitory activity is essential for nervous system function. Neuroactive substances like nicotine can disrupt this balance and hyperactivate dopaminergic circuits, with combined effects on behavior and neural activity. We use *Drosophila* larvae to examine nicotine-induced, linked behavioral changes over multiple time scales by integrating high-resolution locomotor analysis with genetic and pharmacological manipulations. Acute nicotine exposure produces concentration-dependent hyperactivity. Manipulations of the dopaminergic system establish dopamine as the main mediator for motor responses, and further exploration establishes the *γ* lobe of the mushroom body as a key site for nicotine integration. Experiments with long-term and repetitive nicotine exposure suggest sustained circuit excitability. Finally, nicotine exposure history induces nicotine preference, highlighting experience-dependent plasticity contributing to addiction-like behaviors. These results help establish *Drosophila* larvae as a model organism to elucidate how neuroactive substances reconfigure neural circuits and behavior.

## Introduction

### Background

Organisms experience an array of environmental contexts that may affect different neural circuits to elicit context-dependent levels of activity. The balance of excitatory and inhibitory neural activity across circuits is essential for organisms to be able to respond efficiently to their surroundings to ensure survival and fitness. Neuronal activity influences neuron morphology, synaptic strength, and circuit connectivity. Circumstances such as disease states or exposure to neuroactive chemicals can lead to chronic activity in neurons and neural circuits, and therefore behavior. Given the importance of balanced neural activity, it is useful to study sources of chronic *imbalance*, such as chemical stimulants.

The long-term use of stimulants like nicotine, cocaine, and amphetamines is known to produce maladaptive behaviors through mechanisms that alter neurotransmission(***Laviolette and vaan der Koor (2004); Farzam et al. (2022); Wolf et al. (2004); Benowitz (2009); Tiwari et al. (2020); Alasmari (2020); Konova et al. (2015)***). Dopamine (DA) is a common neurotransmitter by which these drugs act on the nervous system and form drug dependence (***Akalal et al. (2006); Bainton et al. (2000); Kalivas and Stewa (1991); Faraone (2018); Chvilicek et al. (2020)***); dopamine is a highly conserved catecholamine neurotransmitter that has neuromodulatory roles in learning and memory, sensory processing, and motor control (***Yamamoto and Vernier (2011); Pérez-Fernández et al. (2021); Ryczko and Dubuc (2017)***). How stimulants cause dysfunctional signaling in the DA neurotransmitter system and how this impacts neural circuit activity and behavior are poorly understood in a single system. A comprehensive understanding of the mechanisms by which drugs cause disruptive neurotransmission and how this contributes to behavioral changes is crucial in order to treat addiction. Vertebrates typically possess millions or billions of neurons, so a simpler model organism should make characterizing the interplay between neural circuits and behavior more tractable.

*Drosophila* larvae have fewer than 10, 000 neurons, an extensive body of data on their nervous system structure, and genetic tools to measure and control neural activity (***Le Bras (2020); Scheffer et al. (2020)***). Their short life cycle, small, translucent body, and simple and slow behavioral repertoire are conducive to a deeper study of the impacts of nicotine on locomotor behavior and the role of the DA neurotransmitter system.

In this paper, we leverage the *Drosophila* larva to characterize the acute locomotor effects of nicotine, determine the trajectory of recovery, and assess the persistence of response after repetitive exposure. We also identify the neural populations and brain regions that engage with nicotine. We examine experience-dependent plasticity by asking how prior exposure affects navigation on a nicotine gradient. Together, these experiments combine high-resolution behavioral analysis with genetic and pharmacological manipulations to characterize how acute and repetitive nicotine exposure reshape locomotion and navigation by reconfiguring dopaminergic neural circuit dynamics. This work connects neural circuit perturbation to its manifestations in behavior, and highlights *Drosophila* as a potential model for probing addiction-related processes.

### *Drosophila* larva locomotion

The biological basis for behavior is the computational and integrative processes of the central nervous system (CNS) (***Schmidt (1989)***). *Drosophila* larvae perform locomotive behavior continually to seek out favorable environments, especially food, because sufficient nutrition is critical for growth and metamorphosis (***Clark et al. (2018)***). Their locomotion, produced by central pattern generators (CPGs) located in the ventral nerve cord (VNC) (***Mantziaris et al. (2020)***) consists of crawling and other movements such as turns, head swings, pauses, rolling, and burrowing (***Clark et al. (2018); Lahiri et al. (2011)***). Some properties of the neural circuits that govern locomotion have been characterized, including the specific sensory and motor neurons that innervate the larval body walls carrying input and output from the CNS (***Clark et al. (2018); Hunter et al. (2021); Lahiri et al. (2011); Song et al. (2007); Zarin et al. (2019)***). Forward and backward crawling, classified as axial locomotion, occur when synchronized contractions of segmented muscle groups propagate from posterior to anterior segments in peristaltic waves (***Kohsaka et al. (2019); Caldwell et al. (2003)***). The CPG’s output drives the rhythmic contractions that form a peristaltic wave, which can occur in the absence of descending inputs (***Caldwell et al. (2003); Kohsaka (2023)***). Thus, the role of the VNC in motor output is relatively well understood. Sensory processing from the peripheral nervous system (PNS) is relayed to motor circuits in the CNS to fine tune motor output and contribute to fast reflexes. Less understood is the role of higher neurons in the larval brain lobes, but it has been shown that these higher neurons or descending inputs can directly modulate behavior (***Clark et al. (2018); Lee and Doe (2021)***). Both the CNS and PNS are thus important in understanding locomotor behavior.

Crawling is the most prominent and well understood behavior, where *runs* move the animals forward, but larvae perform other locomotive tasks that are essential for proper negotiation of their environment. *Turns* underlie decision-making, allowing a larvae to actively respond to sensory experiences(***Gomez-Marin and Louis (2012); Luo et al. (2010)***). While turns may be executed autonomously, sensory inputs from the environment are important for proper behavioral responses; these inputs are processed by the brain, leading to biases in locomotor output that contribute to directed navigation. Similarly, *head swings* assist in altering movement, where larvae perform asymmetric contractions of anterior segments in order to actively sense their immediate environment (***Gomez-Marin and Louis (2012); Lahiri et al. (2011)***). Other common behaviors include burrowing, used to facilitate feeding, and rolling, used to escape nociceptive stimuli (***Kim et al. (2017); Tracey et al. (2003)***). Larvae can perform other behaviors such as bending, hunching, and rearing, but these behaviors are less common and less well characterized (***Gowda et al. (2021)***).

These components of larval locomotion serve as building blocks for complex exploratory and navigational behavior. Reorientation maneuvers are performed according to the sensory integration of stimuli detected during exploratory movements like head swings and crawling (***Gomez-Marin and Lou (2012); Klein et al. (2015)***). Larvae can actively modulate turn characteristics to bias their navigation towards favorable environments (***Lahiri et al. (2011); Gomez-Marin and Louis (2012); Jovanic et al. (2019); Klein et al. (2015***, 2017)). They exhibit navigational behaviors including chemotaxis, thermotaxis, phototaxis and anemotaxis, amongst other responses to outside stimuli (***Kim et al. (2017); Gomez-Marin and Louis (2012); Klein et al. (2015); Yu et al. (2023); Gowda et al. (2021); Jovanic et al. (2019)***).

This rich locomotor repertoire provides quantitative metrics for the effects of neuroactive substances on behavior. In this paper, we characterize the set of larval locomotor behaviors in larvae to understand how nicotine perturbs motor circuits and characterize dopamine’s role in producing maladaptive locomotor patterns.

### *Drosophila* and dopamine signaling

Dopamine (DA) is one of the main neuromodulatory neurotransmitters in the central brain and is linked to modulation of locomotion across many taxa (***Yamamoto and Seto (2014); Joshua et al. (2009); Masoud et al. (2015); Mustard et al. (2010); Oliveri and Levin (2019)***), including *Drosophila*. Dopamine manipulation in the CNS has been shown to impact central motor pattern activity and directed navigation, cause aberrant crawling behavior, hyperactivity, and modulation of specific motor programs like turning (***Cooper and Neckameyer (1999); Fox et al. (2006); Jovanic et al. (2019); Neckameyer (1998); Selcho et al. (2009); Suster et al. (2003)***). Characterization of dopaminergic clusters and their functional roles have identified two main regions in the adult fly and larval brain linked to motor output: the central complex and the mushroom bodies (MB), a region rich in dopaminergic innervation and crucial for reward, sensory processing, and learning (***Scheffer et al. (2020); Xie et al. (2018); Hartenstein et al. (2017); Martin et al. (1998); Silva et al. (2020); Varnam et al. (1996)***). However, the specific dopaminergic neurons responsible for modulating locomotor behavior are not known. Understanding the role of DA as a potential neuromodulator and identifying the key neurons involved would provide insight into how DA regulates behavioral changes.

### *Drosophila*, dopamine, and neuroactive chemicals

Exposure to stimulants, such as nicotine, often alters behavior, often mediated by dopamine. Mammals, including rodents, exposed to stimulants display altered locomotor activity, develop conditioned preferences, and show changes in cognitive performance; however, studying the underlying biological mechanisms for these responses proves difficult due to the complexity of these organisms (***Kalivas and Stewart (1991); Lowenstein and Velazquez-Ulloa (2018); Philyaw et al. (2022); Domino (2001); Lobina et al. (2011); Fowler and Kenny (2011)***). Some invertebrate models show similar responses to stimulants, suggesting that the underlying mechanisms mediating these effects are evolutionarily conserved. Bees with overstimulated circuits associated with rewards can develop a preference for low levels of nicotine, and demonstrate behavioral effects such as excessive grooming (***Williamson et al. (2014); Baracchi et al. (2017)***). *C. elegans* have demonstrated hyperactive locomotor responses to nicotine, changes in behavior due to repetitive exposure, and the ability to associate cues as a reward when paired with nicotine (***Polli et al. (2015); Feng et al. (2006); Salim et al. (2024)***). Thus, the conservation of nicotine-induced behavioral responses across diverse taxa underscores the value of invertebrate models as systems for dissecting the biological mechanisms associated with stimulant-driven hyperactivation.

Adult and larval *Drosophila* can adapt to their environment, with many behaviors found to exhibit plasticity and the ability to enter persistent states as a product of repeated neural activation and changes in the balance of excitation and inhibition (***Kallman et al. (2015); Jung et al. (2020); Asahina et al. (2009)***). Adult flies exhibit hyperactive locomotion, increased grooming, and hyperkinesia in response to stimulants, including nicotine, consistent with overactive dopamine signaling (***Philyaw et al. (2022); Bainton et al. (2000); Andretic et al. (2008); Chvilicek et al. (2020); McClung and Hirsh (1998); Landayan and Wolf (2015); Hou et al. (2004); Zhang et al. (2016); Nall et al. (2016); Blosser et al. (2020)***). One study described the direct synaptic connection between cholinergic and dopaminergic neurons in the brain, through which nicotine could drive dopaminergic activity (***Zhang et al. (2016)***). Most studies of neuroactive chemical response in *Drosophila* have focused on short-term exposure in adult flies (***Kong et al. (2010); Bainton et al. (2000); Hou et al. (2004)***), which does not address the impact of prolonged activation, often associated with consumption of nicotine, and the mechanisms by which neural circuits, connectivity, and behavior may change. Larvae also respond to stimulants, becoming hyperactive when exposed to amphetamines and methylphenidates, likely a dopamine-dependent response (***Pizzo et al. (2013)***). We would expect nicotine to have a hyperactivating effect at some range of concentrations. In addition, nicotine administered directly to the larval CNS excites sensory-motor circuits and stimulates sustained dopamine release in the VNC, but the work was not tied to behavioral analysis (***Malloy et al. (2019); Pyakurel et al. (2018)***).

We build on this previous *Drosophila* work by addressing three key questions: 1) In what ways does nicotine-driven hyperactivaton modulate behavior? 2) What roles do neural circuits play in these responses? 3) What are the mechanisms of drug dependence and addiction-like plasticity? We employ freely crawling *Drosophila* larvae, pairing fine-grained locomotor analysis with neural perturbations. We find that nicotine changes the behavioral repertoire in a concentrationdependent manner, shifting some, but not all, features essential for navigation and exploration; this response reveals key nicotine targets in the context of the motor program. Both pharmacological and genetic manipulation experiments implicate dopamine neurons and the mushroom body, specifically the *γ* lobe, potentially linking neuromodulation to intact behavior. On the scale of hours, repeated exposure continuously drives sustained excitability. In addition, repetitive exposure leads to changes in navigation on a nicotine gradient, suggesting a plastic response within the important neural populations.

These findings should lay the groundwork to establish the *Drosophila* larva as a model to investigate how hyperactivation through neuroactive substances reshapes neural circuit activity and induces changes in behavior. Through detailed high-resolution behavioral characterization coupled with targeted genetic manipulations, we reveal dopaminergic mechanisms underlying stimulant effects and provide a foundation for dissecting the neural basis of drug-induced hyperactivity and its behavioral consequences.

## Results

### Strong nicotine exposure reshapes the behavioral repertoire

We first sought to determine the effect of relatively strong nicotine doses on larval behavior. An ethogram shows the fractional usage of numerous behaviors (Fig. 1A). Following a single exposure to nicotine solution, larvae were tracked over time. Control (water-exposed) larvae demonstrate a typical behavioral profile consisting largely of forward crawling, interspersed with head swings and turns, along with infrequent resting. The behaviors deployed remained consistent across the examined time points (Fig. 1B) and were consistent with previously reported observations (***Gowda et al. (2021); Kohsaka et al. (2014); Lahiri et al. (2011)***).

**Figure 1.**
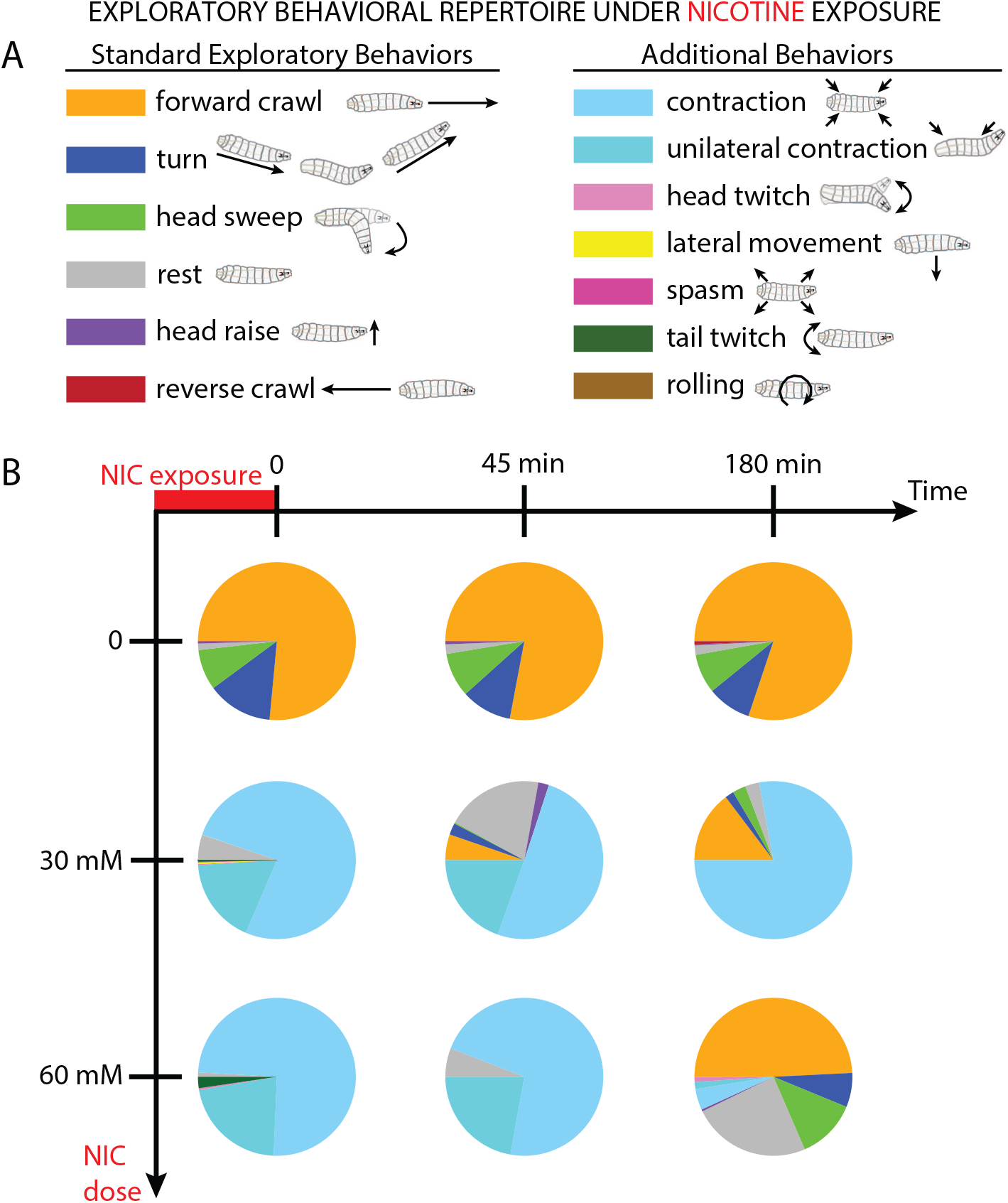
Nicotine exposure alters the behavioral profile of *Drosophila* larvae. **A** Legend and schematics for the set of observed locomotor and postural behaviors. **B** Fractional use of the set of behaviors following nicotine exposure. Between 5 and 7 wild type Canton-S (CS) larvae were used for each condition.

In contrast, administration of nicotine (either 30 or 60 mM) dramatically shifted larval crawling behaviors. Typical activities like crawling, head swinging, and turning were suppressed, replaced by stationary or erratic movements like body contractions, unilateral contractions, and tail or head twitching (Fig. 1B). Larvae exposed to 30 mM nicotine partially recovered 45 min later, resuming more forward crawling, and after 180 min turning and head swinging returned. Recovery from a 60 mM nicotine exposure took more time, with typical locomotor patterns resuming after 180 min. These results demonstrate that nicotine can strongly affect which behaviors larvae deploy, and that larvae can recover from the effects. However, it should be more illuminating to examine more modest doses that modulate typical behaviors, rather than those that render larvae mostly motionless.

### Moderate nicotine exposure alters multiple locomotor features

We focused on lower levels of nicotine exposure in order to find pharmacologically relevant doses that produce strong changes in locomotor parameters, while avoiding toxic doses that largely disrupt locomotion. Larvae were exposed to nicotine solution (or a water control), rinsed, placed on a behavior stage that recorded their locomotion, then subsequent analysis extracted numerous properties (Fig. 2 A,B). We used crawling speed as the primary indicator of hyperactivity, varied the nicotine exposure level, and built the dose-response curve seen in Fig. 2C. Larvae exposed to lower doses from 0 up to 1.125 mM show a dose-dependent increase in crawl speed consistent with the generally expected stimulatory effect of nicotine. We observed the strongest effect at 1.125 mM, a roughly 50% increase in crawl speed. Higher doses reduced crawling speed, which fell well below baseline at ≥ 3 mM, and nearly all larvae ceased crawling by 30 mM. This set of responses is consistent with the previously reported biphasic effect of nicotine on adult *Drosophila* (***Malloy et al. (2019); Wolf and Heberlein (2003); Chvilicek et al. (2020)***). The peak response dosage of 1.125 mM was used in all subsequent experiments in this work.

**Figure 2.**
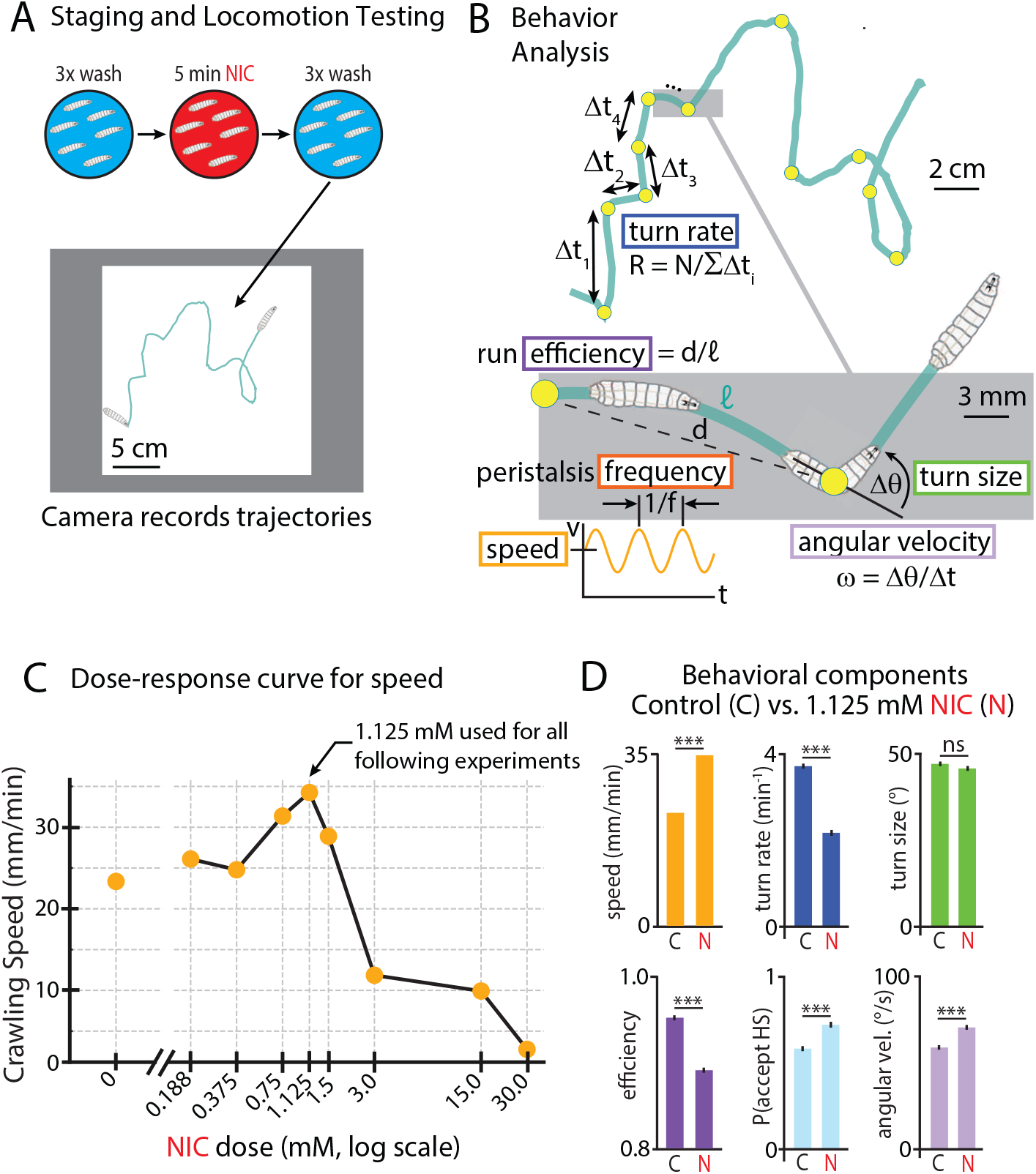
Nicotine affects multiple behaviors in larval locomotion. **(A)** Larvae were rinsed (blue), exposed to a nicotine solution (red), rinsed again, and moved to an agar gel platform for behavioral recording (example track shown in green). **(B)** Extracted behavioral metrics. Example track comprised of an alternating sequence of runs (of duration Δ*t*_*i*_) and turns (yellow circles), which occur at rate *γ*. Properties of runs (efficiency *d*/𝓁, speed ⟨*v*⟩, peristalsis frequency *f*) and turns (size Δθ, acceptance probability *P*, angular velocity ω) are extracted from recorded video data. **(C)** Dose-response curve for moderate nicotine concentrations, with 1.125 mM producing the greatest speed increase. 2191 larva tracks were analyzed, between 88 and 394 for each NIC dose. **(D)** Nicotine changes specific behavioral components. Bar graphs compare six behaviors with and without NIC exposure. Error bars are s.e.m. *** indicates *p <* 0.001, ns indicates *p >* 0.05, Student’s t-test. 394 control (C) tracks of Canton-S larvae were analyzed, and 155 NIC larvae.

Several other behavioral parameters (illustrated in Fig. 2B) also change in response to nicotine exposure (Fig. 2D). Nicotine-treated larvae turn less frequently and their paths are less direct. During turns, they swing their heads more quickly and accept head swing directions more often, while the angle of their turns remains the same. Notably, all of these behavioral changes, except for reduced run efficiency, cause larvae to diffuse away from their starting points more quickly: they move faster, and spend less time performing turns where their forward locomotion is paused. These results suggest that nicotine significantly impacts numerous specific components of behavior, potentially reflective of stimulatory effects impinging on motor circuits.

To further understand the behavioral effects of nicotine, we zoomed in to examine the peristaltic muscle movements of the larvae, and asked how the undulating patterns change with overall speed. We extracted the average speed and persitalsis frequency for thousands of individual runs. Without nicotine, histograms of speed and peristalsis frequency (Fig. 3A) show wide distributions of both properties. There is a negative correlation between the two behavioral features, where faster crawling is driven by slower peristalsis contractions. With nicotine exposure, the distributions shift to higher speeds and lower peristalsis frequencies, with an even stronger negative correlation between the two (Fig. 3B). Side-by-side comparison of two representative trajectories, each with speed and frequency very close to its population’s average values, illustrate the effects of nicotine on crawling motion (Fig. 3C). The nicotine-exposed larva moves farther and crawls faster, but with a longer period peristaltic wave. Overall we find a counterintuitive result where faster crawling does not arise from faster peristalsis, and this holds both within control and nicotineexposed populations and across the two populations.

**Figure 3.**
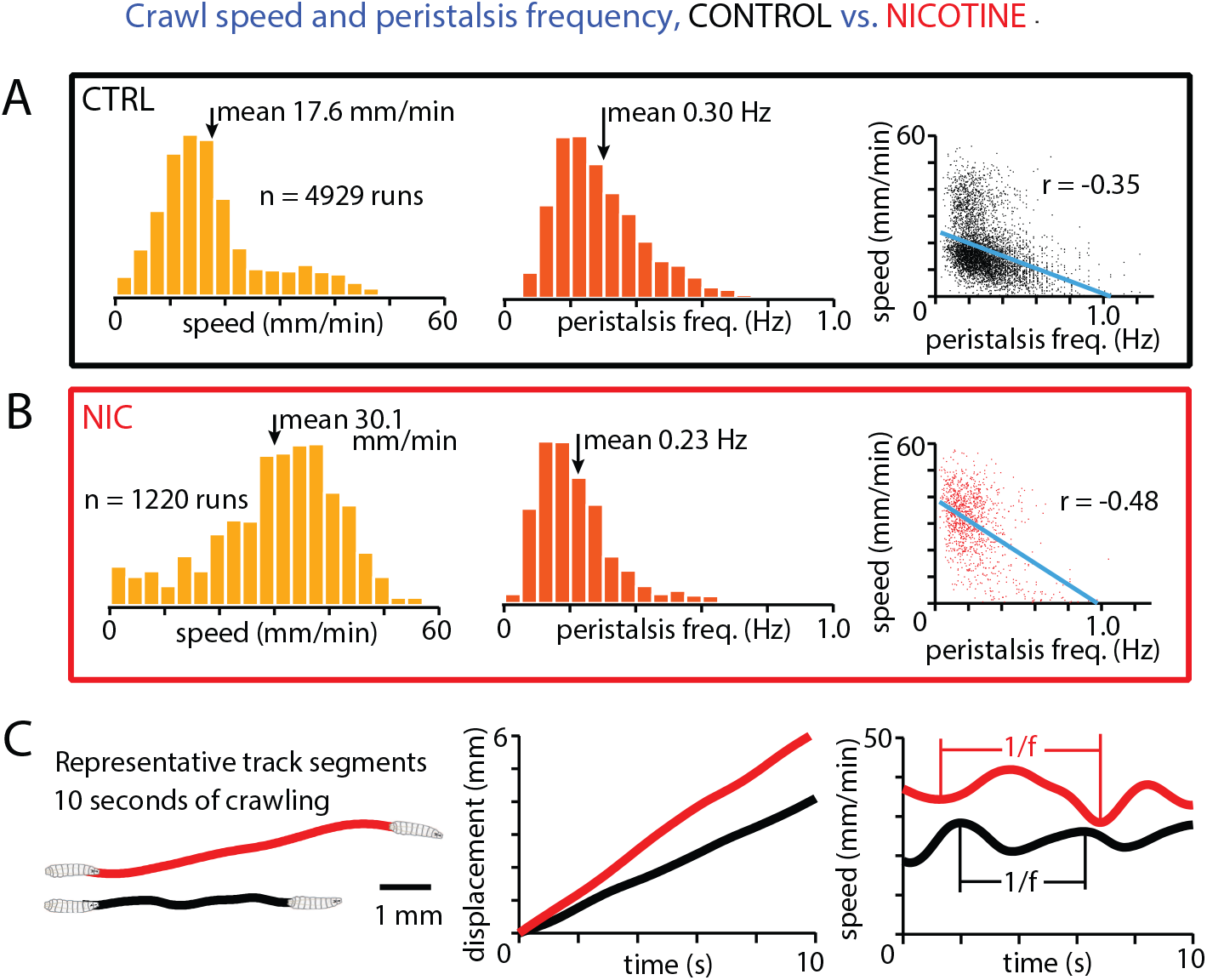
Speed and peristalsis frequency linked and modulated by nicotine. **(A)** Histograms of the average speed (left) and peristalsis frequency (middle) from 4929 runs, and a scatter plot (right) of each run showing the correlation (blue line) between the behavioral features. **(B)** Same information as displayed in panel A, but for larae exposed to 1.125 mM nicotine solution. **(C)** One representative track each for control and nicotine conditions. Each track has a speed and peristalsis frequency extremely close to the corresponding distribution mean. Left: representative tracks plotted as starting from the same point. Middle: displacement of the larva over time for each tracks. Right: speed over time for each track, with vertical lines indicating adjacent peaks or troughs in the sinusoidal pattern. Canton-S larvae from 7 experiments for each condition (394 control tracks, 155 nicotine tracks) were analyzed, the same animals as Fig. 2D.

### Nicotine-induced hyperactivity depends on dopamine

Having established the significant and specific effects of nicotine exposure on larva behavior, we sought to determine what neurotransmitter systems and neural populations were involved in the behavioral responses to stimulants like nicotine. Dopamine is a likely candidate, being linked to mediation of reward responses, responses to neuroactive chemicals, and locomotion. The larval CNS has roughly 120 dopaminergic neurons, including four primary dopaminergic clusters: DL1, DL2, DM1, and pPAM (Fig. 4A,B).

**Figure 4.**
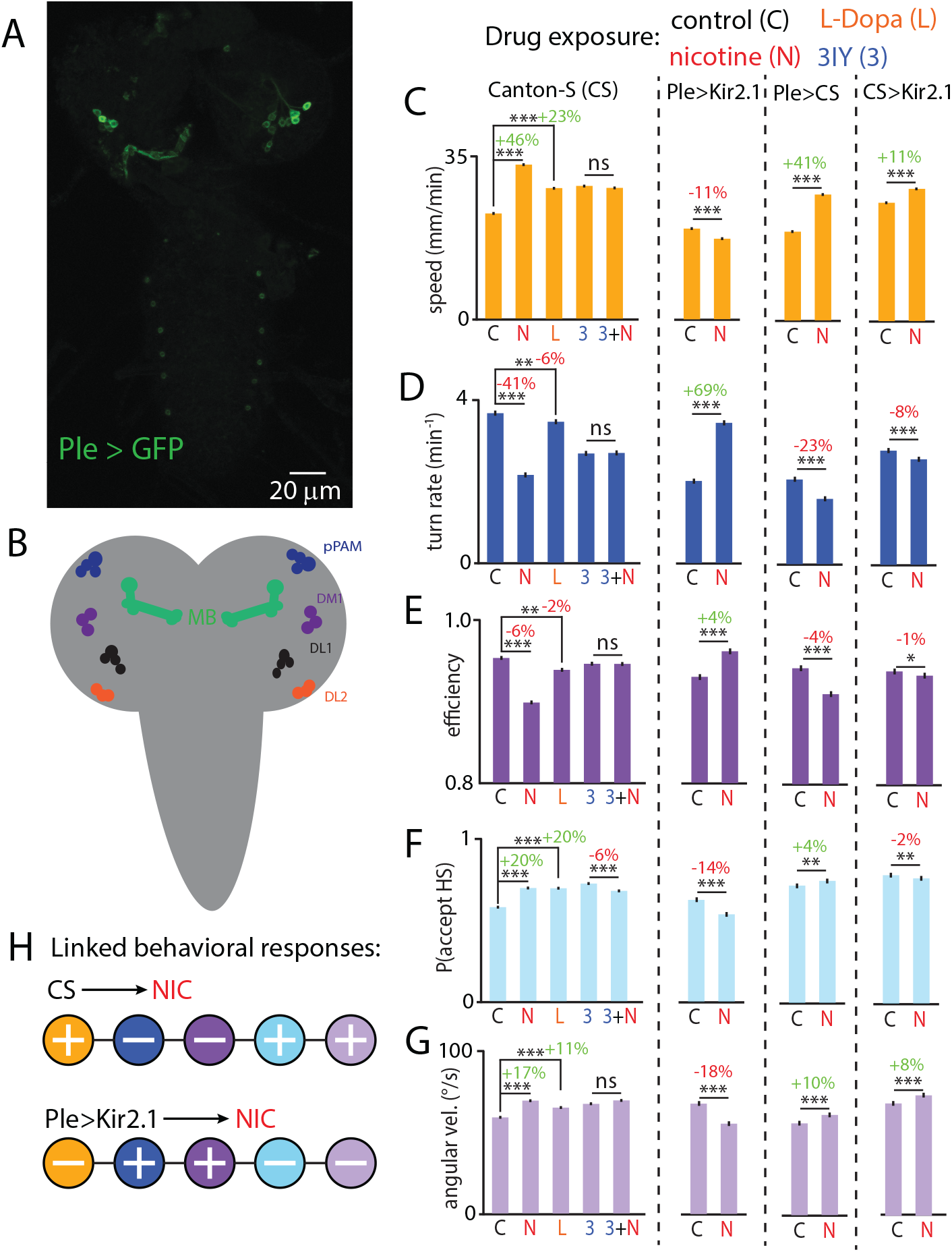
Dopamine is a key component of nicotine-driven hyperactivity. **(A)** Confocal micrograph of an acutely dissected PLE-GAL4 > UAS-Kir2.1-eGFP brain, labeling dopamine neurons in the brain lobes. **(B)** Schematic of larval brain lobes and the ventral nerve cord (VNC) in gray with neural clusters of the dopaminergic system: dorsalateral 1, 2 (DL1,DL2), dorsomedial 1 (DM1), and primary protocerebral anterior medial (pPAM), and the mushroom body (MB), important in the dopaminergic system. **(C-G)** Left: Bar graphs showing the behavioral impact of four pharmacological treatments compared to control (C): nicotine (N), L-Dopa (L), 3IY (3) and nicotine and 3IY together (3+N). Treatments were administered as described in Methods and Materials. Right: Bar graphs showing the behavioral impact of genetic silencing of dopaminergic neurons on the response to nicotine exposure. Different strains are separated by dashed lines. Percent changes are indicated above lines (red for decreases, green for increases). **H** Schematic summary of the directional changes in behavioral components with and without silenced dopamine neurons. Between 150 and 400 larvae were used for each experimental condition (2381 total). Error bars are s.e.m. *** indicates *p <* 0.001, ** indicates *p <* 0.01, * indicates *p <* 0.05, ns indicates *p >* 0.05, Student’s t-test.

We combined pharmacology and genetic manipulation of dopaminergic neurons to uncover the role of dopamine in the behavioral response to nicotine. As shown above (Fig. 2D), nicotine significantly increased crawling speed, suppressed turning rate, reduced run efficiency, and affected head swing acceptance and velocity, also seen in the leftmost bars of Fig. 4C,D,E. Direct application of L-Dopa, a precursor to dopamine, partially phenocopied all five of these responses, by amounts less than or equal to the 1.125 mM nicotine exposure. Treatment with 3IY, a pathway inhibitor for dopamine synthesis, prevented effects of nicotine across locomotion (Fig. 4C-G), where head swing acceptance decreased slightly, but no other behavior significantly changed. Larvae treated with 3IY did demonstrate changes to locomotion compared to untreated controls, so comparisons of nicotine response were made relative to the 3IY baseline behavior. Thus, nicotine reliably drives changes across at least five behaviors, with L-Dopa eliciting similar but weaker effects, while 3IY effectively blocked the nicotine-induced behavioral responses.

We also suppressed dopamine neurons with genetic silencing, by expressing the potassium inward rectifier Kir2.1 in dopaminergic neurons. These Ple*>*Kir2.1 larvae still changed their behavior when exposed to nicotine, but with the *opposite* valence as the original wild type controls and both heterozygote controls. Speed decreased while turn rate and efficiency increased; head swing acceptance diminished along with head swing angular velocity, all with strong statistical significance (Fig. 4C-G, and summarized schematically in Fig. 4H). This reversed response could emerge from the absence of dopamine neural activity, where the actions of nicotine on non-dopaminergic pathways become more prominent, emphasizing dopamine’s role as a key modulator of nicotine-induced behavior. The results here do demonstrate that the hyperactive effect of nicotine on this set of behaviors depends on dopamine signaling.

### Nicotine-induced hyperactivity operates in mushroom body circuitry

Having determined that nicotine exposure alters specific behavioral features, and that dopaminergic neurons play a key role in this process, we next asked which specific brain regions mediate the nicotine-driven changes in larval locomotion.

The mushroom body (MB) in the *Drosophila* larva is a neuropil that receives extensive dopaminergic innervation (***Tanaka et al. (2008); Bang et al. (2011); Rohwedder et al. (2016); Saumweber et al. (2018); Aso et al. (2014)***). Several patterns of cells make up the MB lobes: the *a*,, *p*,, and *γ*-born neurons. These lobes contain intrinsic mushroom body neurons, the Kenyon cells, along with a complex neuropil composed of axons and dendrites that receive input from diverse sources. Specifically the *γ*-born neurons bifurcate to form the vertical and medial lobes in the larval brain, while the *a*, and *p*, neurons develop later (***Pauls et al. (2010); Qi et al. (2025); Truman et al. (2023); Hartenstein et al. (2017); Silva et al. (2020)***).

To probe the MB’s contribution to nicotine-induced changes in crawling, we used OK107-GAL4 driving Kir2.1 to silence the entire MB, and 201Y-GAL4 to silence the *γ*-born neurons (Fig. 5A). As shown above (Figs. 2 and 4), larvae exposed to nicotine crawl faster and less efficiently, turn less frequently, swing their heads faster, and are more likely to accept head swings. The silencing of the neurons using either OK107-GAL4 or 201Y-GAL4 significantly attenuated the nicotine-driven response in these metrics (Fig. 5B-F): all five behavioral changes were either dramatically reduced, abolished, or even mildly reversed. Heterozygote controls maintained their nicotine responsiveness. Since neural silencing with either driver attenuates the nicotine responses, the *γ* lobe appears to be the key brain region involved in the process. But since the behavioral changes are not entirely eliminated, it is possible that neurons outside the *γ* lobe are necessary for the complete response to nicotine.

**Figure 5.**
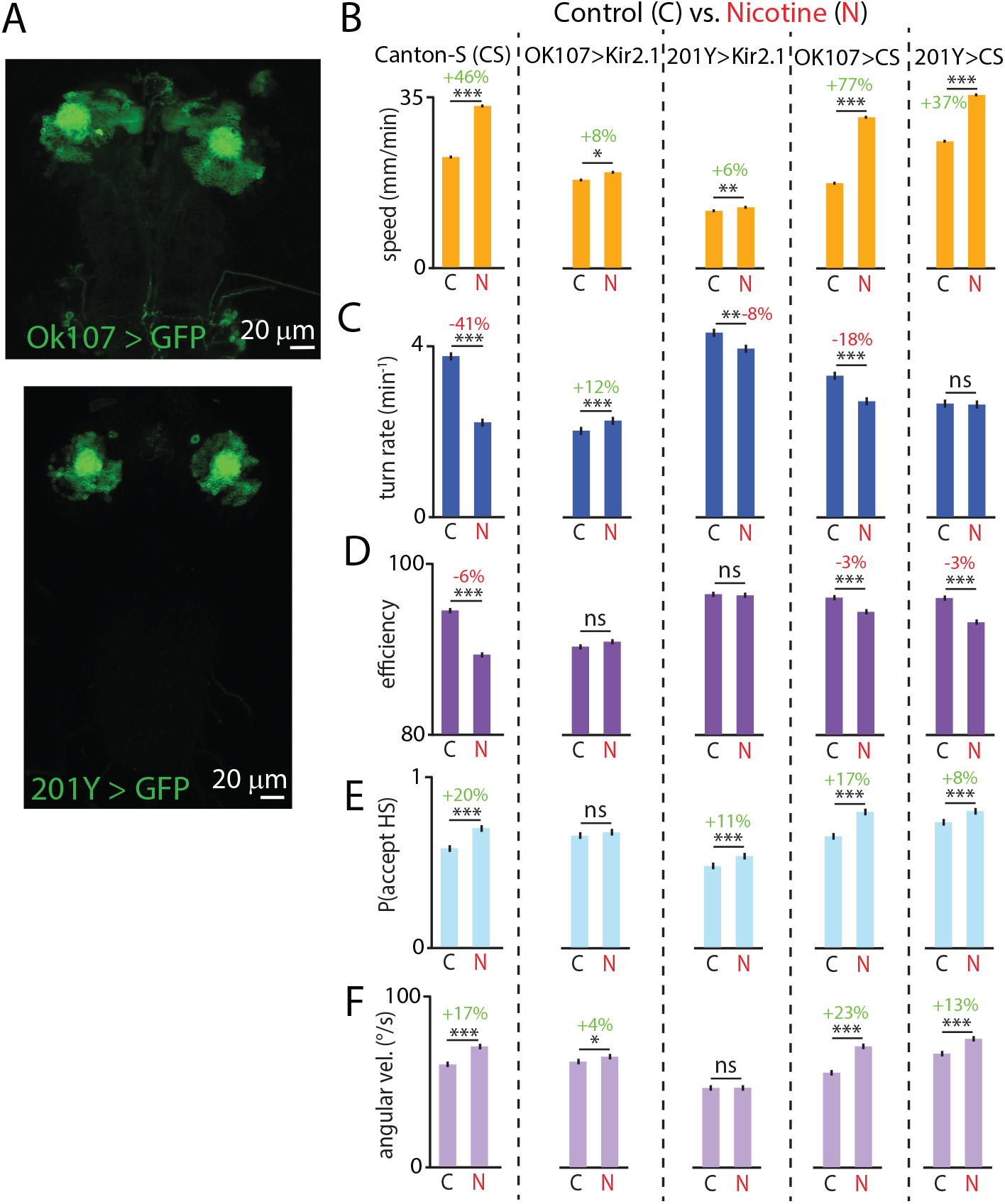
The mushroom body (MB) is an important brain region involved in nicotine-driven hyperactivity.**(A)** Confocal micrographs of an acutely dissected OK107-GAL4 > UAS-Kir2.1-eGFP brain (top) and a 201Y-GAL4 > UAS-Kir2.1-eGFP brain (bottom), labeling the entire MB and the *γ* lobe, respectively. **(B-F)** Bar graphs for five behavioral features showing the impact of MB manipulation on the response to nicotine (N) treatment. Different strains separated by dashed lines. Percent changes are indicated above lines (red for decreases, green for increases). Between 150 and 400 larvae were used for each experimental condition (2372 total). Error bars are s.e.m. *** indicates *p <* 0.001, ** indicates *p <* 0.01, * indicates *p <* 0.05, ns indicates *p >* 0.05, Student’s t-test.

### Longer-term nicotine exposure induces sustained behavioral effects

Having observed that moderate nicotine exposure induces significant changes in multiple behavioral features (Fig. 2), and that larvae recover even from strong nicotine exposure (Fig. 1), we examined nicotine response over longer time scales.

We started by characterizing the recovery of larval behavior (specifically crawling speed) following exposure to nicotine (Fig. 6A). After a single 5-min exposure, crawling speed showed an immediate increase and then declined over time with a time constant of approximately 30 min. By 90 min, crawling speed returned to the level of untreated larvae.

**Figure 6.**
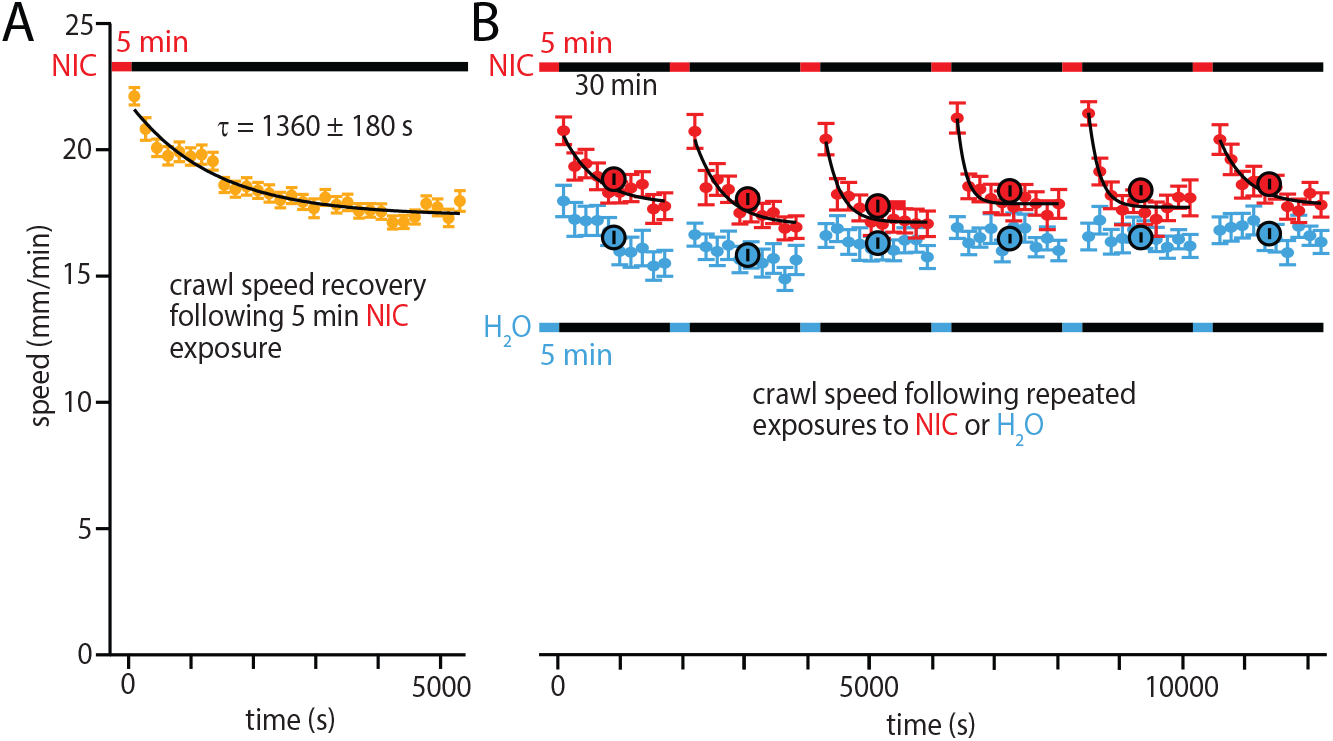
Recovery from single and repeated nicotine exposures. Horizontal bars show exposure protocols, with red bar and text indicating nicotine, blue water control, and black the free crawling during recording. **A** Single nicotine exposure recovery. Crawling speed (circles) indicate the mean speed in 3-min time windows. Black curve is an exponential decay function fit (*y*(*t*) = *y*_0_ + *Ae*^−*t*/*τ*^). 88 larvae were recorded. **B** Repeated nicotine exposure recovery. Solid blue (control) and red (nicotine) circles indicate mean speeds in 3-min time windows, and larger black-outlined circles the mean of the whole 30 min segment between exposures. Black curves are exponential fits for nicotine-exposed larvae (control data is not exponential). 62 larvae used for control experiments and 54 larvae for nicotine experiments. Canton-S larvae were used in all experiments.

Repeated stimulant use often drives changes in neural circuit activity and disrupts homeostatic regulation to produce profound and lasting impacts on behavior (***Lowenstein and Velazquez-Ulloa (2018); Philyaw et al. (2022); Wolf et al. (2004); Ping and Tsunoda (2012); Kalivas and Stewart (1991)***). We used our fly larva model system to quantify the effects of repeated nicotine exposure (Fig. 6B), submerging groups of larvae in nicotine solution six times while measuring crawling behavior between exposures. Repetitive exposure to nicotine led to an increase in crawl speed compared to water-exposed controls. All six nicotine exposures caused a rapid increase in speed, followed by steady decline over 30 min, similar to the single-exposure response in Fig. 6A. This cycle of hyperactivity persisted across all six exposures, suggesting that habituation does not occur within the three-hour time frame. We note that speed did also change modestly for the control group, an effect that declined after the first hour of the experimental assay.

### Repeated nicotine exposure induces navigation on a nicotine gradient

Behavioral response to nicotine is sustained over the hours-long times we have investigated so far. A logical follow-up question is whether prior nicotine exposure could lead to addiction-like states, such a preference for the (normally neutral or aversive) compound. To answer this question, we tested larval movement on a spatial nicotine gradient after prior exposure. Figure 7A,B demonstrates a two-solution casting method (based on the salt gradients from ***Luo et al. (2014)***) that produces a linear gradient of nicotine concentration in an agar gel after 48 hrs.

**Figure 7.**
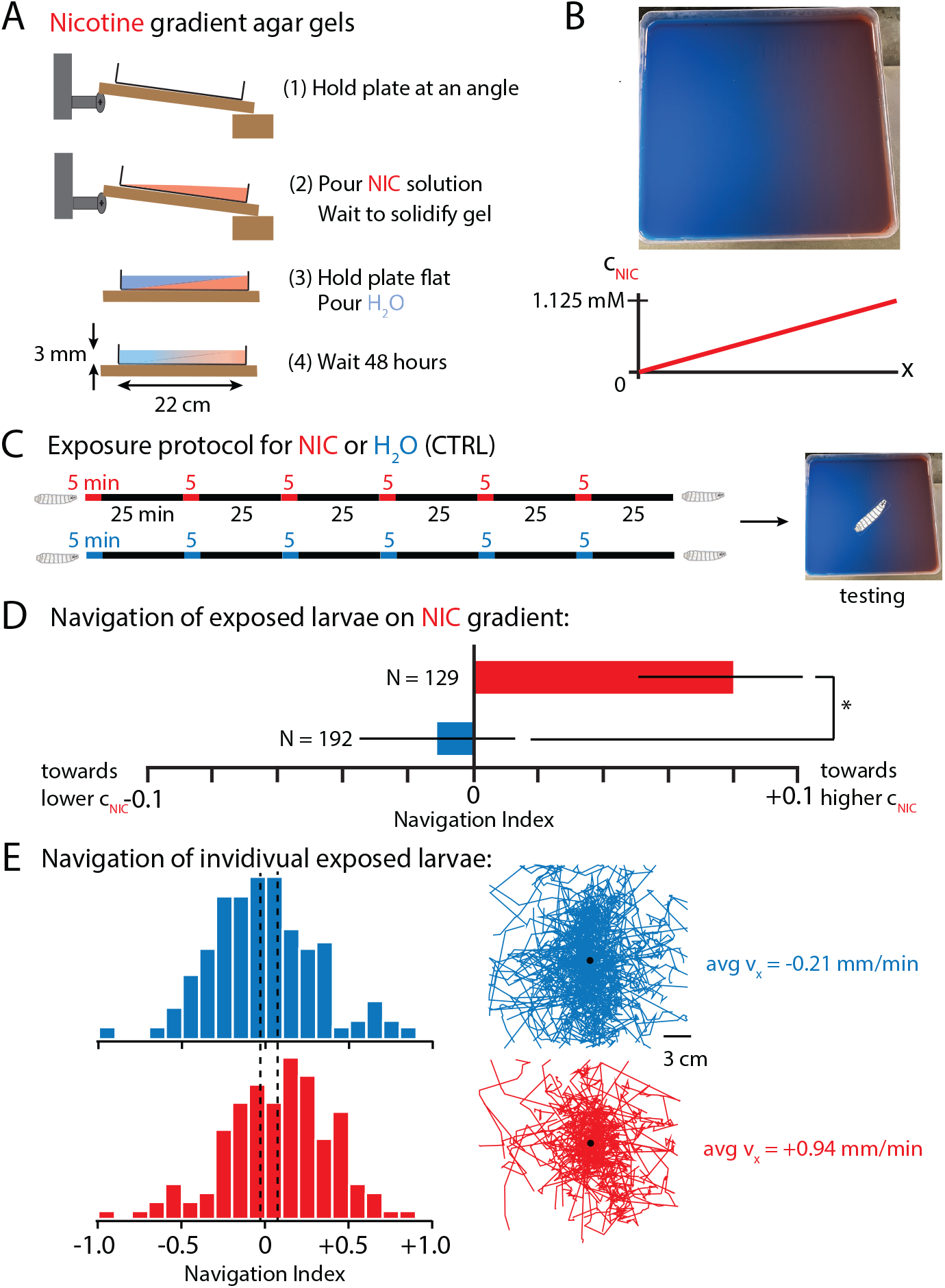
Larvae navigate towards nicotine after repeated exposure. **A** Protocol for making nicotine gradient gels (blue dye solution is water and red the nicotine). **B** Photograph of the gradient gel, and a schematic graph of its linear horizontal nicotine gradient from 0 to 1.125 mM. **C** Protocol for the nicotine exposure prior to testing. Larvae are submerged in water (blue), or submerged in 1.125 mM nicotine (red), or freely moving on plain agar gel (black). **D** Net larval movement along the nicotine gradient summarized by the mean navigation index, each larva has an individual index *NI* = ⟨*v*_*x*_⟩/⟨*v*⟩. Positive *NI* indicates navigation towards nicotine. *N* indicates the number of larvae tested per condition. **E** Navigation indexes of individual larvae. Left: Distribution of *NI* for control (blue) or nicotine-exposed (red) larvae. Vertical dashed lines indicate the mean *NI* of the populations. Right: Crawling trajectories for all experiments and the average *x* component of velocity. * indicates *p <* 0.05, Student’s t-test. Canton-S larvae were used in these experiments.

Larvae were given repeated 5 min exposures to either nicotine or water control over a 3 hr period with 30 minute segments of resting (Fig. 7C), then placed on the nicotine gradient to test navigation. Larvae with prior nicotine exposure exhibited statistically significant positive navigation up the gradient towards higher concentrations of nicotine. Water-treated control larvae navigated slightly down the gradient. This navigation difference was quantified with both average navigation index (*NI*) (Fig. 7D) and with distributions of individual larvae *NI* and overall population drift velocity (Fig. 7E). This demonstrates that prior nicotine experience can change larval navigation towards nicotine, potentially a promising start to using the animal as a model for chemical addiction.

## Discussion

In this work we aimed to characterize the effects of nicotine-driven hyperactivation on *Drosophila* larvae over both short (minutes) and longer (hours) time scales at the behavior level, and identify the relevant components at the neural circuit and neurotransmitter levels. Previous studies of nicotine response in flies, with both adults (***Bainton et al. (2000); Hou et al. (2004); Zhang et al. (2016); Ren et al. (2012)***) and larvae (***Pyakurel et al. (2018); Malloy et al. (2019)***), have provided insight into the process, but used coarser behavior metrics over narrower time scales. To our knowledge, a systematic analysis of the locomotor response to acute and repetitive exposures of hyperactive chemicals has not been performed, and the regions of the brain that mediate these responses have not been identified.

### Nicotine drives hyperactivity at low doses and replaces behaviors at higher doses

By tabulating the physical movements of fly larvae following exposure to strong nicotine concentrations (Fig. 1), we found that overstimulation replaced normal crawling motions (runs and turns) with stationary and erratic behaviors like contractions and twitching. Most likely, these nicotine doses disrupted normal circuit dynamics and impaired coordination (***Giniatullin et al. (2005); Feduccia et al. (2012); Sobkowiak et al. (2011)***). Importantly, larvae did recover over time, on the order of hours. We expect this recovery profile to depend on the nicotine dosage, but the limited number of animals we tested in this type of experiment with manual labeling of behaviors prevent us from making specific claims about higher dose recovery. For example, the fact that recovery appears faster at 60 mM nicotine exposure than at 30 mM could be an artifact of high variance in behavior in general, specifically when high doses involve saturation of motor circuits. An important aspect of understanding how neuroactive chemicals modulate behavior and neural circuitry function is characterizing the behavioral profiles through time. These broader results for high nicotine doses clearly show that nicotine can dramatically alter behavior profiles, but we spent the remainder of this work on lower doses while characterizing stimulant effects on standard locomotor features like the properties of forward crawling and turning.

### Crawl speed and peristalsis frequency are negatively correlated

At lower concentrations, nicotine stimulates movement, notably crawling speed, but depresses speed at higher concentrations (Fig. 2C), a biphasic effect consistent with drug studies in other invertebrate models (***Malloy et al. (2019); Polli et al. (2015); Sobkowiak et al. (2011)***). This effect may correspond to different states of the underlying target receptors: from 0 to 1.125 mM we observe hyperactivity indicating an effect on nicotinic acetylcholine receptors (nAChRs) through cholinergic pathways; above this level, speed decreases, potentially caused by disruption of neural activity in cholinergic pathways that result in motor control dysfunction. Our working dosage of 1.125 mM nicotine maximally drives hyperactivity, reflected in faster forward movement and other changes to crawling and turning (Fig. 2D). Using this particular concentration allowed us to more readily observe behavioral changes during pharmacological and genetic manipulations.

In examining the connection between overall forward motion and muscle movement patterns, we uncovered a counterintuitive result that peristalsis frequency actually decreases in conjunction with faster crawling, both within larval populations and when comparing nicotine-exposed larvae to the control group (Fig. 3). That is, induced hyperactivity, or faster crawling in general, is not equivalent to making every physical movement happen faster.

Previous studies have demonstrated that speed modulation arises from internal circuit dynamics regulating the activity of the central pattern generators (CPGs), including the A27-GDL feedforward circuit that directly controls forward locomotion, activity in command-like neurons within the VNC (AcN, A01j, A02j), or descending inputs from brain lobes (***Zarin et al. (2019); Carreira-Rosario et al. (2018); Clark et al. (2018); Kohsaka et al. (2014)***). Mechanistically, the changes we see may arise from nicotine-driven excitation of cholinergic neurons in these circuits that may alter the duration of muscle contractions or alter the neural timing, allowing changes in cadence and stride length (***Malloy et al. (2019); Kohsaka et al. (2014); Sun et al. (2022); Zhao et al. (2024)***). Overall these behavioral changes may be a consequence of the dual action of nicotine through the CNS and ventral nerve cord (VNC) of the larvae. Acetylcholine, as a major excitatory neurotransmitter, is prominently expressed in the central brain and in the VNC, where it may bind to interneurons and motor neurons to promote excitability. Consistent with this idea, ***Malloy et al. (2019***) found that nicotine drives excitation across all levels of the nervous system, from sensory input and processing in the CNS to motor neurons.

### The role of dopamine in hyperactivating locomotion

Nicotine is known to facilitate dopamine release in mammals (***Benowitz (2009); Tiwari et al. (2020)***), and a study using fast scan cyclic voltammetry showed that in the larval VNC, nAChR stimulation led to sustained dopamine release, possibly through dopaminergic modulatory interneurons (***Gowda et al. (2021); Pyakurel et al. (2018)***). Dopamine also functions as a key neuromodulator of the motor program in *Drosophila* through higher order control centers in the brain lobe. Manipulation of dopamine levels has been shown to affect basal activity, although there are conflicting findings as to whether it increases or decreases locomotion (***Yamamoto and Seto (2014)***). However, increasing dopamine signaling with diverse neuroactive chemicals has been shown to increase locomotion in both adult flies and larvae (***Pizzo et al. (2013); Kong et al. (2010); Bainton et al. (2000)***). Therefore, it is possible that nicotine targets the larval VNC and the crawling motor program through dopaminergic potentiation of the CPGs or changes in the central brain region. The CPGs are crucial for generating locomotive behaviors like crawling, turning, and head sweeping, and could lead to the hyperactive changes in behavior seen in Fig. 2 (***Gowda et al. (2021); Hunter et al. (2021)***).

Dopamine has also been found to play a direct role in the propagation of forward crawling. Application of high concentrations of dopamine promote forward locomotion, and optogenetic manipulation experiments match our observations of increased peristaltic wave duration (***MacLeod et al. (2025)***). This supports the idea that dopamine may be similarly involved in the results demonstrated in Fig. 3.

Dopamine may also act through top down modulation of locomotion, a function attributable to the mushroom body (***Silva et al. (2020); Martin and Krantz (2014); Mustard et al. (2010)***). Alternatively, these two mechanisms could act in parallel across the different dopaminergic circuits, potentially explaining the distinct effects we see across behavior (Fig. 2D). This dual action model would be consistent with findings that dopamine functions in the planning and execution of exploratory behaviors and thus will have differential effects across locomotion (***Silva et al. (2020); Neckameyer (1996)***).

We directly tested the role of dopamine in the hyperactive behavioral changes in Fig. 4, finding that reducing global dopamine levels in the CNS with 3IY or silencing dopaminergic neurons with Kir2.1 prevented the hyperactive effect of nicotine across all behavioral parameters. The administration of L-DOPA, a precursor to dopamine, phenocopies the effects nicotine expsoure, suggesting that dopamine causes the behavioral changes. The magnitudes of changes observed with L-DOPA are weaker, possibly because the dose we administered did not match the amount of stimulation, or because, unlike nicotine, L-DOPA is an endogenous molecule that can be metabolized by the organism efficiently (***Pyakurel et al. (2018)***). Our results here are consistent with previous work in adult flies where dopamine is similarly involved in the behavioral response to nicotine (***Bainton et al. (2000); Zhang et al. (2016)***), and we have built on prior work in larvae by connecting nicotine exposure to dopaminergic circuits to behavioral changes.

We also note that 3IY exposure by itself appears to induce behavioral changes that are similar to nicotine, although weaker for the dose we used. These effects are consistent with what previous observations showing reduced turning rate and stronger head swings (***Thoener et al. (2021)***). This may be due to the role of dopamine as a neuromodulator of locomotor behavior, rather than simply activating or inhibiting locomotion.

### Hyperactivation changes a linked set of behaviors

We found that silencing dopaminergic neurons in larvae resulted in a valence switch in the response to nicotine across all five behaviors highlighted (Fig. 4H). This switch may reflect the action of nicotine through non-dopaminergic pathways. Nicotine activates nAChR’s that are widely present throughout the *Drosophila* nervous system, including kenyon cells in the MB and motor, projection, and inter neurons in the larval CNS (***Rosenthal and Yuan (2021); Korona et al. (2022); Wu et al. (2017)***). Effects on these pathways might be secondary to the hyperactivation seen with dopamine release, but with the dopamine neurons silent the effects of nicotine may switch locomotion patterns to the opposite valence. Alternatively, the silencing of dopaminergic neurons across development may lead to complex homeostatic changes that alter how nicotine affects circuits. Future studies would benefit from more precise temporal control over dopaminergic activity during the relevant time windows, as suggested by subsequent experiments (Fig. 6).

Apart from questions about dopaminergic pathways, the linkage between the five behaviors (speed, run efficiency, turn rate, and head swing speed and acceptance) is interesting in itself. These five features (or six if we include peristaltic frequency, although this was not measured in most experiments) might tend to vary together in larval crawling behavior. Our own prior work (***Evans et al. (2023)***) has noted a strong negative correlation between speed and turn rate within and across larval instars. Except for the relatively small change in crawling efficiency, the valence of the other behaviors all act to increase the rate that larvae move away from a location. Crawling faster with fewer and shorter turns increases the diffusion constant in a random walk, for example. Circuit signals could hyper- or hypo-activate behavior by manipulating all behaviors at once. Understanding how these behaviors are linked together at the circuit level will be important in future studies of locomotor circuits in the brain, in this organism and likely others as well.

### The role of the mushroom body in nicotine-driven hyperactivity

The observation that dopamine plays a role in nicotine-driven hyperactivity led us to investigate the mushroom body, due to its rich dopaminergic innervation, as a potentially key brain region. We found that the MB, particularly the *γ* lobe, is essential for the response to nicotine across all behaviors (Fig. 5). This finding fits with the DL1 and pPAM clusters innervating the medial and vertical lobe made from *γ*-born neurons. Additionally, the *γ* lobe is known to express Dop1R receptors and kenyon cells within the lobe respond to nicotine. It is possible the *γ* lobe serves as an important relay center for nicotine excitation that is propagated through downstream neurons to impact locomotor circuits (***Silva et al. (2020); Leyton et al. (2014)***). However, some behaviors such as turn rate or head swing acceptance still change in response to nicotine. This may be due to dopamine neurons outside the MB remaining responsive to nicotine. Further studies targeting specific dopaminergic neurons or small clusters would help refine our understanding of the neural circuit, by characterizing the roles the neurons play in behavioral response.

### Temporal dynamics of recovery from hyperactivation

Establishing the recovery timeline to nicotine exposure(s) should help reveal the underlying circuit dynamics and responsiveness. We found that the hyperactiving effect on crawling speed decays in approximately 30 min, suggesting that short-term nicotine effects are reversible (Fig. 6A).

We also found little evidence of habituation to nicotine over a few hours of repeated exposure. Immediately after each nicotine solution submersion, the crawling speed jumped to the same value, establishing a persistent, stable nicotine responsiveness (Fig. 6B). We note that the speed appeared to decrease more quickly starting with the third exposure, but further studies with larger sample sizes would be needed to confirm this.

The recovery dynamics imply that the relevant neural circuits, within this time frame, do not experience receptor inactivation and continue to display hyper-responsiveness. Extending these studies would further explore habituation and tolerance, especially if extended past 24 hrs, and would aid in understanding circuit adaptations and the associated behavioral changes.

### The *Drosophila* larva as a potential model for chemical addiction

Acute behavioral plasticity can also occur within shorter time frames (***Feng et al. (2006)***), so we followed the repeated exposure experiments with testing of navigation performance on a nicotine gradient, to see if prior exposure induces a preference for the substance.

We found that control larvae with no prior exposure have a slightly negative navigation index (*NI*) (Fig. 7D), but those with prior nicotine exposure crawled in the opposite direction towards higher nicotine concentrations.

This *NI* magnitudes are small compared to some robust larval behaviors like thermotaxis (***Evans et al. (2023); Klein et al. (2015)***), but comparable to modest appetitive chemotaxis navigation (***Gomez-Marin et al. (2011)***). Notably, these experiments did not involve associative conditioning to induce a learned response, but rather the navigation bias we observe reflects how exposure alone dramatically alters the larval response to the gradient.

The results suggest that, given an exposure history, larva can develop a preference for nicotine and that underlying circuits may already be demonstrating plasticity in their ability to alter larval navigation. A plausible target for this is the dopaminergic clusters that innvervate the MB, a critical region for reward-related behaviors (***Devineni and Scaplen (2022); Bjordal et al. (2014); Lowenstein and Velazquez-Ulloa (2018); Schumann et al. (2021); Eschbach et al. (2020)***). This result is consistent with what has been found in *C. elegans* where the worms could be conditioned to prefer nicotine (***Salim et al. (2024)***). We have demonstrated what could be interpreted as “addictive” behavior, but a more systematic study that optimizes the exposure protocol is warranted. A more efficient method of inducing nicotine navigation preference would be valuable for testing response to other types of stimulants, with a quantifiable addiction metric in hand.

A useful next step would be to manipulate the neurons and brain regions indicated in our study here to test the effect on nicotine gradient navigation, potentially with thermogenetic or optogenetics tools. The experiments from Fig. 6 help establish the appropriate timeline. Additionally, given that this work characterized how locomotion varies in response to nicotine, future studies could benefit from applying a fine-scale behavioral analysis approach to expanding our understanding of the possible molecular and circuit basis for the observed changes to navigation.

### Conclusions

Overall, this paper establishes *Drosophila* larvae as a strong model organism to study how hyperactivity reshapes neural circuits and behavior. By carefully dissecting the behavioral effect of nicotine and identifying dopamine and the mushroom body as critical mediators, we provide a framework through which to dissect the neural mechanisms of drug-induced hyperactivity and potentially addiction related behaviors.

## Methods and Materials

### Fly strains and husbandry

The strains used in this paper are wild type Canton-S (CS, BDSC#64349), UAS-Kir 2.1 (BDSC#6596), Ple-Gal4 (BDSC#8848), OK107-Gal4 (BDSC#854), and 201Y-Gal4 (BDSC#4440). Fly stocks were maintained at room temperature in vials or bottles with cornmeal and molasses food. Larvae used for experiments were raised on grape juice agar plates (1.5%) with a paste made from deionized (DI) water and inactive yeast (0.35-0.45 g). Grape juice plates were changed every 24 hours. Second instar larvae (48-72 hr after egg laying) were used in all experiments in this work, and were selected based on size, mouth hook, and spiracle development.

### Larva preparation

Larvae were rinsed in DI water and were either exposed for 5 min to 400 *µ*l of DI water or to aqueous nicotine (TCL) solutions. The larvae were rinsed again with DI water and moved to a plain agar gel transfer plate prior to behavioral experiments.

Some larvae were fed the dopamine-synthesis inhibitor 3-Iodo-L-tyrosine (3IY) (VWR) or the dopamine precursor 3,4-dihydroxyphenylalanine (L-DOPA) (VWR), both diluted with DI water and stored in closed bottles at 4 °C and wrapped in Al foil to prevent light exposure. Yeast plates were made by adding 400 *µ*l of 5 mM L-DOPA or 32 mM 3IY to yeast with a drop of blue food coloring (Loraan Oils) to identify the larvae that ingested the paste. Larvae were left to feed on the yeast paste for 48 hours and were sorted as second instars.

To make 32 mM 3IY, the compound was slowly added to the DI water as the solution was heated to approximately 30 °C. This solution was made fresh each day, to avoid precipitation of 3IY, then incorporated into the yeast paste. Larvae were left to feed for 24 hrs and selected at second instar. Ingestion was tracked through blue food coloring in their gut, with any larvae lacking blue coloring excluded. The larvae were then exposed to DI water or aqueous nicotine solutions.

The behavioral repertoire of larvae exposed to high nicotine concentrations was characterized by manual observation of recorded activity. Following exposure to 0 (control), 30 or 60 mM nicotine, larvae were transferred to a 60 mm Petri dish of agar and their free behavior was video recorded at 0, 45 and 180 min with a camera (AmScope) affixed to a dissection microscope. Control larvae were individually recorded to be able to track the entirety of their locomotor movement. Larvae exposed to nicotine moved much less and were recorded as groups.

### Behavioral crawling arenas

A temperature-controlled platform (***Evans et al. (2023)***), set to 25 °C, was used for observing larval locomotory behavior after treatment. Agar gels for the platform were 22×22 cm squares with a thickness of 3.5 mm. The gel composition was 2.5% wt./vol. agar (MilliporeSigma), with 0.75% wt./vol. charcoal (Fisher Scientific) to enhance image contrast. The agar gels were placed atop the platform along the midline and acted as crawling substrates for recording larval behavior.

For the nicotine gradient experiments, we made an agar gradient gel similar to the protocol used in ***Luo et al. (2014***) for salt gradients. The first agar solution (1.125 mM nicotine) was poured at an incline (1.13°) forming a wedge volume, allowed to cool, returned to a flat position, then the DI water was added. Nicotine was indicated by red dye and DI water indicated by blue dye. The gel was allowed to set for 48 hours while diffusion of nicotine created a linear gradient along one direction.

### Image acquisition

The behavior platform was surrounded by red LED strips (620 nm, not visible to larvae) arranged in a square perimeter pointing inward to provide dark-field illumination. A CCD camera (Basler) mounted above the behavioral platform with an 8-mm lens (Computar) recorded crawling larvae continuously at 15 frames per second.

Long term and recovery videos were recorded on a robotic platform, a separate system described in ***Yu et al. (2023***). The resultant mp4 file was converted to stacks of JPEG files through FFmpeg software and then run through the customary analysis pipeline described below.

### Data analysis

Raw video data was run through the MAGAT Analyzer software pipeline (***Gershow et al. (2012)***) that extracted each larva’s position and body contour at each frame. The resulting tracks of the larvae were segmented into *runs* of continuous forward movement and *turns* that changed the crawling direction. Tracks that were excessively slow or too short were removed except when examining high nicotine doses.

Custom analysis using Matlab and Igor Pro derived key locomotion and navigation metrics: crawling speed, turning rate, turn size, run efficiency, head swing acceptance rate and head swing angular velocity. For experiments with a stimulus gradient we calculated the navigation index (*NI*), a dimensionless summary metric that represents the efficiency of navigation along an axis. This is computed as the average *x*-component of each larva’s velocity during runs divided by the runs’ average speed: *NI* = ⟨*v*_*x*_⟩/⟨*v*⟩.

Analysis for higher nicotine concentrations required additional manual scoring due to toxic effects that rendered a larger proportion of larvae stationary, but not dead (determined by persistent mouth hook movement and contractile behavior). Stationary larvae were then included in the calculation of average crawling speed in this case.

Customized Igor Pro software also extracted the larva’s peristaltic frequency during forward crawling. For each run with duration ≥ 10 s, an interactive tool allowed the user to select adjacent peaks or troughs from (generally sinusoidal) speed vs. time graphs, then the average peristalsis period, frequency, and forward speed were calcuated for each run.

Ethogram data was analyzed using the software BORIS (***Friard and Gamba (2016)***), with manually-logged behavior types and durations for individual larvae.

### Long term and recovery behavior experiments

Long term behavior experiments repeatedly exposed larvae to a nicotine solution or DI water (control) for 5 min at regular intervals. Larvae were placed on a robotic platform (***Yu et al. (2023)***), but without using its robotic nozzle. In our protocol we paused the recording and manually moved larvae from the edge of the arena to the center Larvae crawled freely for six 30-min windows, interrupted by 5-minute exposures. At the end of each 30-min segment, the larvae were removed from the platform, introduced into the solution, washed, and returned to the platform.

During nicotine gradient testing, the larvae were subjected to above protocol, then moved to the nicotine gradient gel, placed along the midline, and allowed to crawl freely for 10 minutes.

Recovery experiments were conducted by exposing larvae one time to aqueous nicotine solution. Larvae were then placed on the robotic platform and recorded for 90 minutes of free crawling. Each animal was repositioned as described above when it approached the edge of the arena.

## Acknowledgments

The authors thank the University of Miami Biology and Physics departments for internal support. We are grateful to Kevin Collins, Julia Dallman, and Vivek Venkatachalam for comments on the manuscript. We thank Jack Kelty for assisting in linear gradient gel production and James Yu for assistance with robot system troubleshooting.

